# Comparing Methods for Deriving the Auditory Brainstem Response to Continuous Speech in Human Listeners

**DOI:** 10.1101/2024.05.30.596679

**Authors:** Tong Shan, Ross K. Maddox

**Affiliations:** Department of Biomedical Engineering, University of Rochester, United States; Department of Neuroscience, University of Rochester, United States; Kresge Hearing Research Institute, Department of Otolaryngology-Head & Neck Surgery, University of Michigan, United States

## Abstract

Several tools have recently been developed to derive the Auditory Brainstem Response (ABR) from continuous natural speech, facilitating investigation into subcortical encoding of speech. These tools rely on deconvolution, which models the subcortical auditory pathway as a linear system, where a nonlinearly processed stimulus is taken as the input (i.e., regressor), the electroencephalogram (EEG) data as the output, and the ABR as the impulse response deconvolved from the recorded EEG and the regressor. In this study, we analyzed EEG recordings from subjects listening to both unaltered natural speech and synthesized “peaky speech.” We compared the derived ABRs using three regressors: the half-wave rectified stimulus (HWR) from Maddox and Lee (2018), the glottal pulse train (GP) from Polonenko and Maddox (2021), and the auditory nerve modeled response (ANM) from Shan et al. (2024). Our evaluation focused on the fidelity, efficiency, and practicality of each method in different scenarios. The results indicate that the ANM regressor for both peaky and unaltered speech and the GP regressor for peaky speech provided the best performance, whereas the HWR regressor demonstrated relatively poorer performance. The findings in this study will guide future research in selecting the most appropriate paradigm for ABR derivation from continuous, naturalistic speech.

## Introduction

Speech is a complex sound encountered daily and plays a fundamental role in human communication. It is thus essential to understand the process through which the human brain translates speech from its basic encoding by the auditory periphery to higher level processing in the cortex. Subcortical structures, particularly the inferior colliculus, have been proven to be critical in this auditory processing chain, notably in the encoding of vowels and processing speech in noise environments (Carney, Li, & McDonough, 2015). The auditory brainstem response (ABR) serves as a key metric for subcortical auditory neuroscience research as well as clinical audiology. Traditionally, the ABR is characterized by a stereotypical evoked potential elicited by brief stimuli such as clicks, tones or chirps (R. F. Burkard, Eggermont, & Don, 2007; Picton, Hillyard, Krausz, & Galambos, 1974) through electroencephalography (EEG) recording. This evoked potential is observed in the first ∼10 milliseconds post-stimulus, consisting of components that reflect different stages of auditory pathway according to their latency. Specifically, Waves I, III and V are of particular interest, corresponding to the responses of the auditory nerve, cochlear nucleus, and inferior colliculus and lateral lemniscus, respectively (Picton et al., 1974).

Expanding upon this foundation, investigations into the brainstem’s response to speech via complex ABR (cABR) have been undertaken (Krizman, Skoe, & Kraus, 2010; Musacchia, Sams, Skoe, & Kraus, 2007; Skoe & Kraus, 2010). These studies demonstrate that short speech vowels elicit a transient onset and a frequency following response (FFR) corresponding to the voiced part. However, the cABR method has limitations in its controversial neural sources (Coffey, Herholz, Chepesiuk, Baillet, & Zatorre, 2016) and potential neural adaptation due to the repetitiveness of the speech stimuli used (i.e., repeated tokens of vowels or syllables).

Recently, studies have developed several methods for detecting the brainstem response to continuous, non-repetitive speech, thus offering a more ecologically valid approach (Bachmann, MacDonald, & Hjortkjær, 2021; Forte, Etard, & Reichenbach, 2017; Kulasingham et al., 2024; Maddox & Lee, 2018; Polonenko & Maddox, 2021b; Shan, Cappelloni, & Maddox, 2024). One such technique involves extracting the fundamental waveform from the speech and cross-correlating the waveform with the EEG signal (Forte et al., 2017). This method yielded a broad peak around 9 ms primarily originating from the inferior colliculus but was lack of fine temporal responses for other components. Another set of studies are based on a deconvolution method that was proposed by Maddox and Lee (2018). The deconvolution method shared a similar concept as the temporal response function (TRF) used for cortical response to natural stimuli (Di Liberto, O’sullivan, & Lalor, 2015; Ding & Simon, 2012; Lalor & Foxe, 2010; Lalor, Power, Reilly, & Foxe, 2009) and provides superior responses to the fundamental waveform-based methods (Bachmann et al., 2021). As depicted in **Figure 1a**, an encoding model was proposed: the stimulus (more specifically, an acoustical feature derived from the stimulus) acted as the input (i.e., regressor, *x* in **Figure 1a**), the recorded EEG signal as the output *y*, and the ABR as the impulse response of a linear system that transforms *x* into *y*.

**Figure 1.**
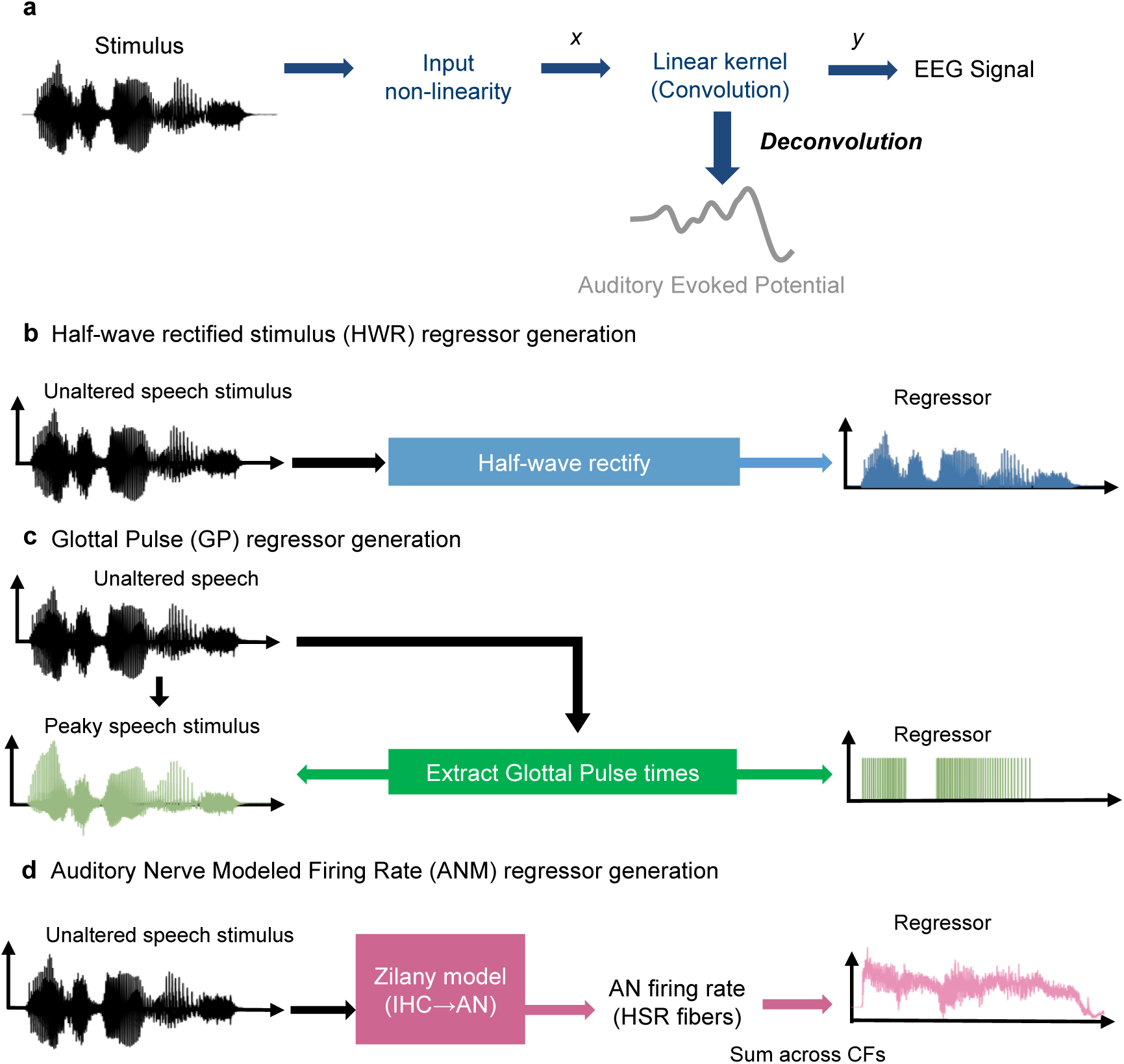
The encoding model using deconvolution method and the regressors that are used. **(a)** The deconvolution paradigm. **(b)** The half-wave rectified stimulus regressor (HWR). **(c)** The peaky speech waveform and the Glottal Pulse train regressor (GP). **(d)** The auditory Nerve Modeled firing rate regressor (ANM). IHC = inner hair cell, AN = auditory nerve, HSR = high spontaneous rate, CF = characteristic frequency.

A series of studies have offered improvements for ABR deconvolution methods. The initial study by Maddox and Lee (2018) utilized half-wave rectification of the stimulus as the regressor (HWR, **Figure 1b**) as a simple simulation of cochlea nonlinearity. It was able to derive the speech ABR with a distinct Wave V that is highly correlated with the click-evoked ABR. Following this, Polonenko and Maddox (2021b) proposed using “peaky speech,” a re-synthesized speech stimulus that was made impulse-like by aligning the phase of the speech harmonics at the time of glottal pulses. The regressor used was a train of impulses placed at the times of the glottal pulses (GP regressor, **Figure 1c**). This method provided distinct earlier ABR components waves I and III in addition to wave V and enabled simultaneous ABR measurements from separate frequency bands. Shan et al. (2024) further extended the deconvolution paradigm by incorporating a detailed computational model that simulates the neural representation of the auditory periphery, converting the stimulus from waveform into auditory nerve modeled response to use as the regressor (ANM, **Figure 1d**), and Kulasingham et al. (2024) compared the ANM to several simpler regressors, finding that a more efficient model provides good responses as long as it recapitulates the adaptation present in the auditory nerve. The ANM method, like the peaky speech with the GP regressor, also yields ABRs with early wave components, and improves the speech ABR’s signal-to-noise ratio (SNR) over the HWR regressor. Moreover, this ANM method is generalizable to other natural sounds, including music.

In this study, we aimed to compare ABR deconvolution using the HWR, GP, and ANM regressors. By examining the fidelity, efficiency, and practicality of each method in different scenarios, we hope to offer guidance on determining the most appropriate approach for deriving ABRs from natural speech as well as other complex sounds for a variety of experimental or clinical uses.

## Results

The data analyzed in this study was obtained from a peaky speech experiment previously conducted by Polonenko and Maddox (2021a). This dataset includes EEG recordings from 22 subjects with normal hearing who were passively listening to an English audiobook under two conditions: unaltered and peaky speech. In the Results section, we present the ABRs derived from the three different regressors and asses their quality using various quantitative metrics. Following this, we introduce a novel approach designed to enable a more equitable comparison of time-domain responses derived from these spectrally different regressors.

### GP and ANM regressors yield quicker and more robust ABR

Using the deconvolution method with the regressors depicted in **Figure 1**, we obtained the ABRs for both unaltered and peaky speech from the three regressors. Illustrated in **Figure 2** are the response derived from HWR (**Figure 2a**), GP (**Figure 2b**) and ANM (**Figure 2c**). By looking at the general waveforms, it is apparent that the GP for peaky speech condition and the ANM for both conditions exhibit better ABR morphology.

**Figure 2.**
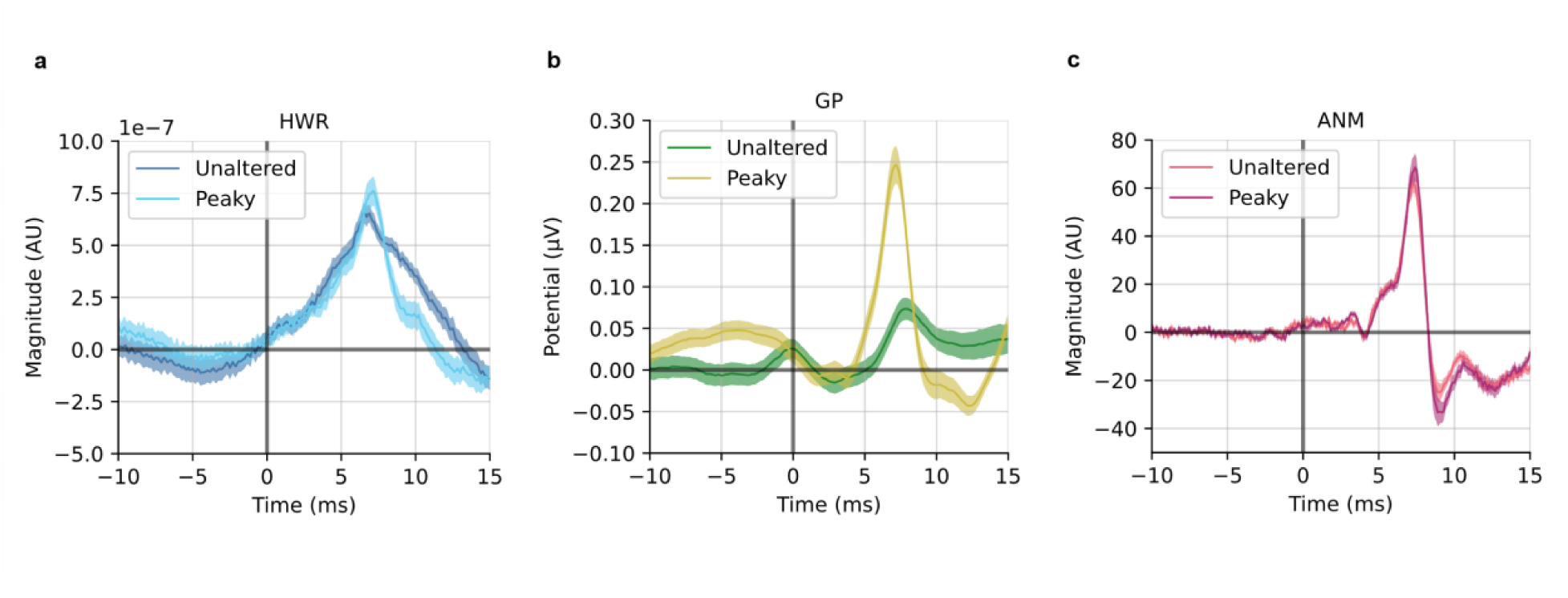
Grand averaged ABR waveforms for unaltered and peaky speech derived from HWR (**a**), GP (**b**) and ANM (**c**) regressor. Shaded area show ± 1 SEM (n = 22).

The ABR derived from HWR shows a broad wave V for both stimulus conditions (**Figure 2a**), consistent with findings from previous studies (Maddox & Lee, 2018; Polonenko & Maddox, 2021b). The ABR derived from GP for peaky speech has a distinct and narrow wave V at around 7.2 ms along with an early component (Wave I) at around 3.2 ms. The GP regressor is not designed for unaltered speech, but since it captures limited acoustical representation for the speech at the glottal pulse time, a much smaller wave V is still observable (**Figure 2b**). Note that we ran this regressor-stimulus combination for completeness, but we did not expect high-quality responses from it. The ANM regressor yields clear ABRs for both unaltered and peaky speech with very similar waveforms and high consistency across subjects. Early components (Wave I and Wave III) were present in the waveforms in addition to Wave V (**Figure 2c**).

We then performed an analysis of the signal-to-noise ratio (SNR) for the ABR waveforms derived from the three regressors within the 0 to 15 ms time window (see Materials and Methods for details of SNR computation). The analysis revealed significant variability in SNR across the regressors for both unaltered and peaky speech (p<0.001; repeated measures ANOVA). The results are shown in **Figure 3a**. For unaltered speech, the ANM regressor demonstrated the highest SNR of 11.81 ± 0.49 (mean ± SEM), which was significantly better than the SNR obtained with HWR, which averaged 4.45 ± 1.07 (p<0.001; two-tailed paired t test, Holm-Bonferroni corrected). As expected, both ANM and HWR showed higher SNR than GP in this condition (p<0.05; two-tailed paired t test, Holm-Bonferroni corrected). For the peaky speech condition, the SNR was the greatest for ANM (12.94 ± 0.53), followed by GP (9.44 ± 0.72) and HWR (5.30 ± 0.73) in that order. Post-hoc pairwise comparison further showed significant differences between each regressor (p<0.001; two-tailed paired t test, Holm-Bonferroni corrected). A similar trend was observed when extending the analysis to a time range of 0 to 30 ms for the derived waveforms (**Figure S1**).

**Figure 3.**
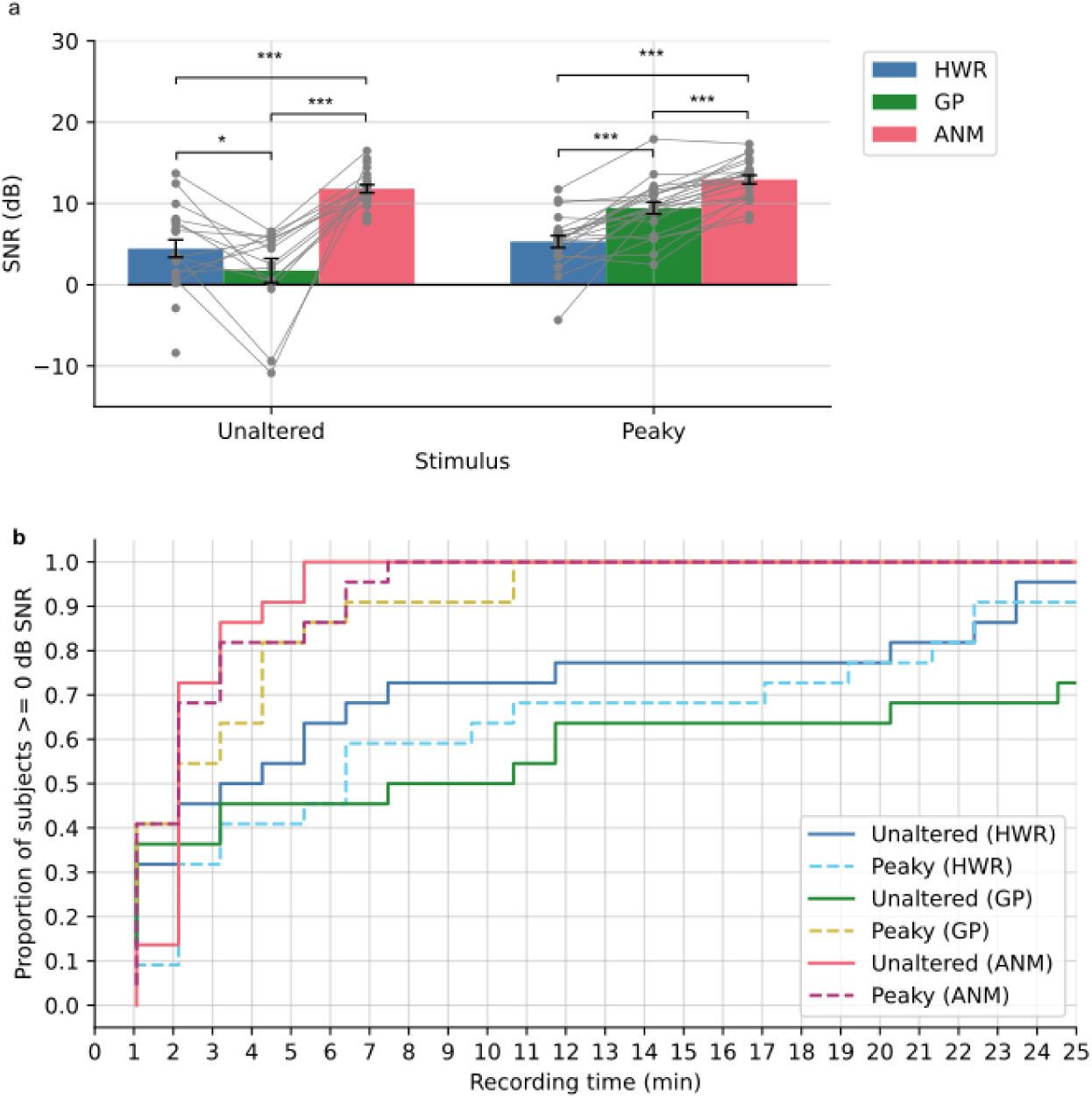
SNR analysis for the derived ABRs. **(a)** The averaged SNR of the ABR for unaltered and peaky speech derived from the three regressors. The bar represents the averaged SNR across subjects and the grey dots with lines are the SNRs for each individual subject. **(b)** The cumulative proportion of subjects that has ABR SNR >= 0 dB as a function of recording time.

We were interested in how efficient each regressor is in deriving a ABR by measuring the time required for subject to achieve a good response with SNR greater than 0 dB. As shown in **Figure 3b**, for unaltered speech, all subjects reach 0 dB SNR within 6.4 minutes when using the ANM regressor. Conversely, when using the HWR and GP regressors for the duration of the 42-minute experiment, only 95% and 86% of subjects reached 0 dB, respectively. With peaky speech, it takes 35.2 minutes for all subjects to reach 0 dB using HWR, while it only takes 11.7 minutes with GP and 8.53 minutes with ANM. These indicate that the ANM is efficient in both conditions, and GP exhibited superior performance in peaky speech. Both the ANM and GP outperformed HWR.

### ABR derived from GP and ANM regressor can better predict EEG

In line with common practices in cortical TRF studies (Crosse, Di Liberto, Bednar, & Lalor, 2016; David, Mesgarani, & Shamma, 2007), we conducted a Pearson correlation analysis to evaluate the accuracy of EEG signal prediction against real EEG recordings utilizing the waveform derived by each regressor. As the previous study (Shan et al., 2024), we initially used the derived waveforms from time range [0, 200] ms as a full kernel, which was then convolved with the regressors to generate the predicted EEG. **Figure 4a** shows the prediction accuracies for unaltered and peaky speech across regressors. There was no significant effect of regressor type in either speech condition (p=0.097 for unaltered and p=0.44 for peaky speech; repeated measures ANOVA). This broadband measure reflected a large portion of signals from cortical activity, indicating a consistent predictive performance across regressors in later component of the auditory potentials.

**Figure 4.**
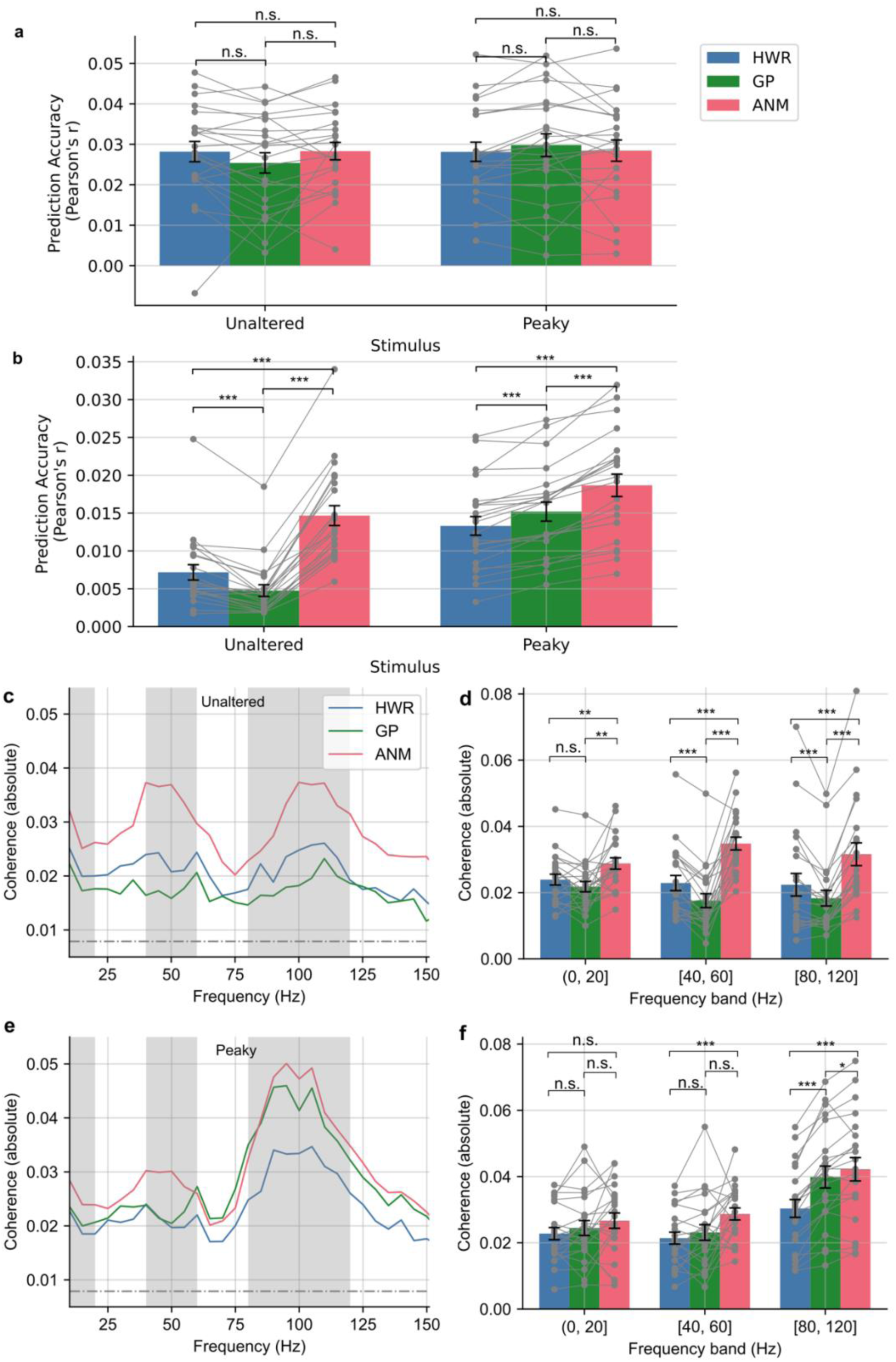
Prediction accuracy as the correlation coefficient and spectral coherence between predicted and real EEG data. **(a)** Broadband correlation coefficient with full kernel (0–200 ms). **(b)** Correlation coefficient of high-pass filtered EEG with subcortical kernel (0–15 ms). The bars are averaged accuracy across subjects with error bars showing ± 1 SEM and the grey dots with lines are for each individual subject. **(c)** The mean absolute value of spectral coherence for unaltered speech. **(d)** The mean coherence in three frequency bands for unaltered speech across subjects. **(e)** The mean absolute value of spectral coherence for peaky speech. **(f)** The mean coherence in three frequency bands for peaky speech across subjects. The dash-dotted lines in **(c)** and **(e)** indicate the noise floor. The shaded grey areas indicate the frequency bands analyzed in **(d)** and **(f)**. The bars in **(d)** and **(f)** are averaged coherence across subjects with error bars showing ± 1 SEM and the grey dots with lines are for each individual subject. (*p<0.05, **p<0.01, ***p<0.001)

We then narrowed our focus to a shorter time range of 0 to 15 ms to emphasize subcortical encoding. In both unaltered and peaky speech, the prediction accuracies were low since the later, slower cortical component of the EEG was not part of the model (but were still present in the signal). There were also no differences among the regressors (p=0.257 and p=0.099, respectively; repeated measures ANOVA).

However, a distinct divergence among regressors emerged upon applying a high-pass filter at 40 Hz to the EEG signals to de-emphasize slower cortical activity, significant for both speech conditions (p<0.001; repeated measures ANOVA; **Figure 4b**). Specifically, in the unaltered speech condition, we again observed that both HWR and ANM demonstrated better accuracy compared to GP (p<0.001; two-tailed paired t test, Holm-Bonferroni corrected). Additionally, ANM exhibited an advantage over HWR (p<0.001; two-tailed paired t test, Holm-Bonferroni corrected). In the peaky speech condition, GP and ANM both outperformed HWR (p<0.001; two-tailed paired t test, Holm-Bonferroni corrected), and ANM also showed significantly better accuracy than GP (p<0.001; two-tailed paired t test, Holm-Bonferroni corrected). Notably, mixed effects linear regression showed that the correlation between the two stimulus conditions were significantly different with peaky speech having higher coefficients (p=0.029 for stimulus condition variable).

In addition to the correlation analysis, we conducted a spectral coherence analysis to evaluate the models’ prediction accuracy across frequency, similar to the approach utilized in Shan et al. (2024). This analysis quantifies the normalized similarity between the predicted and actual EEG data at each frequency, providing detailed insights into model performance on a per-frequency basis (see Materials and Methods for details). **Figure 4c and 4e** highlights the superiority of the ANM regressor over GP and HWR in unaltered speech and the advantage of ANM and GP over HWR in the peaky speech condition. These coherence trends are consistent with the comparative superiority of ANM for unaltered speech and ANM and GP in peaky speech seen with other metrics.

Shan et al. (2024) identified significant advantages of the ANM regressor over HWR particularly in the frequency ranges centered around 50 Hz and 100 Hz. Therefore, we further break down the coherence comparison into three frequency bands: (0, 20] Hz, [40, 60] Hz and [80, 120] Hz (**Figure 4d and 4f**). We then conducted a statistical comparison of the mean coherence from the three frequency bands across the regressors. We found that the ANM regressor outperformed the other two regressors in in all three bands for unaltered speech (p<0.01; two-tailed paired t test, Holm-Bonferroni corrected; **Figure 4d**). In the peaky speech condition, both ANM and GP exhibited superior performance compared to HWR in [80, 120] Hz, and ANM was slightly superior compared to GP (ANM vs HWR, p<0.001; GP vs HWR, p<0.001; ANM vs GP, p=0.02; two-tailed paired t test, Holm-Bonferroni corrected). The ANM regressor was also found to show higher coherence than HWR in [40, 60] Hz band, but not significantly higher than GP (ANM vs HWR, p<0.001; ANM vs GP, p=0.06; GP vs HWR, p=0.22 two-tailed paired t test, Holm-Bonferroni corrected). However, no significant advantage was observed in the low-frequency band (p=0.53; repeated measures ANOVA; **Figure 4f**).

### An Approach for Fair Comparison Using the Phase-only Regressor

While our analysis demonstrated that ANM regressor, especially when combined with peaky speech stimuli, offers an advantage across multiple metrics among the three regressors, it is crucial to acknowledge the inherent spectral differences among the regressors as illustrated in **Figure 5**. The deconvolution process, which includes dividing the Fourier transform of the EEG signal by the Fourier transform of the regressor, highlights the significance of the regressor’s spectrum on the result response spectrum, which effectively acts as a filter, where frequencies with lower amplitude in the regressor are emphasized in the resulting deconvolution. (It should be noted that even if the analysis is done in the time-domain, the same still applies, as the stimulus autocorrelation is accounted for in the time domain.) For example, the ANM regressor (**Figure 5c**) has a decreasing magnitude in higher frequency regions compared to the GP (**Figure 5b**), leading to the ABR derived from ANM containing larger magnitudes in higher frequencies than that derived from the GP.

**Figure 5.**
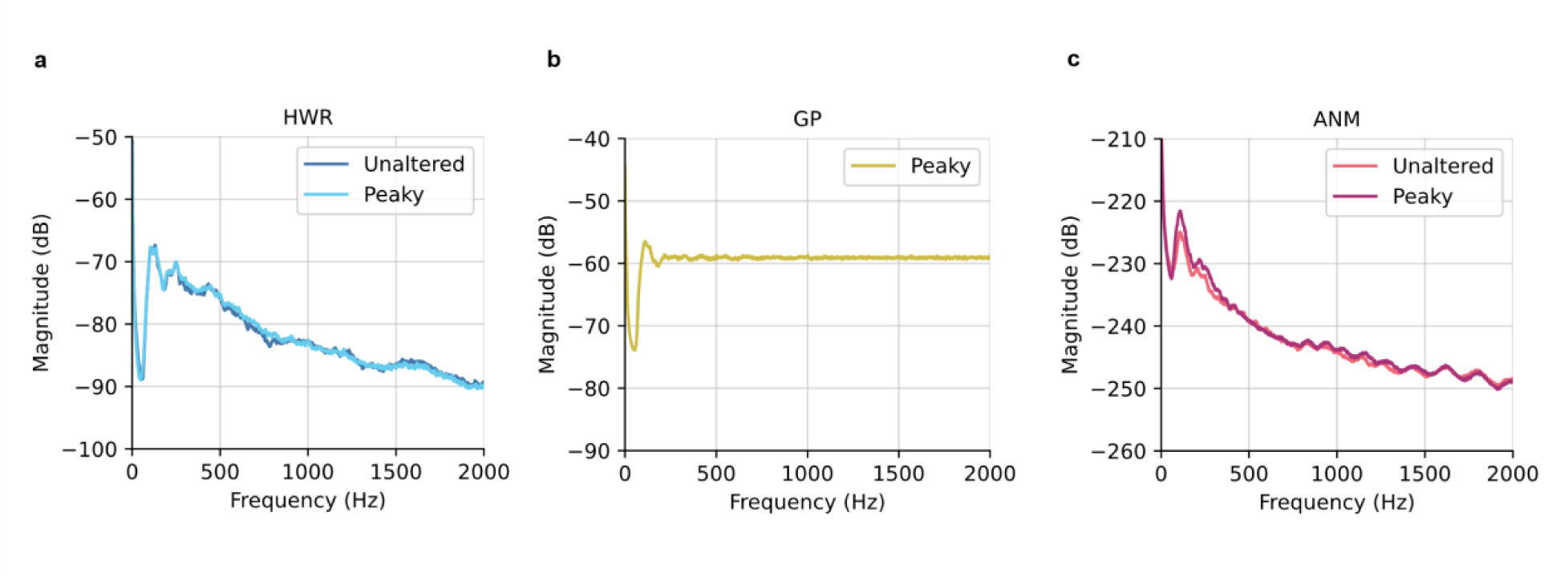
Averaged power density spectrum with Welch estimate for HWR **(a),** GP **(b),** and ANM **(c).**

To account for these spectral differences, we introduced a more equitable comparison method by utilizing a phase-only regressor, where the magnitude of each regressor is set to unity at all frequencies, keeping only the phase to capture the temporal information inside the regressor (see Materials and Methods for details on making phase-only regressor).

The derived response from the three phase-only regressors preserved the characteristic morphology of ABRs, as depicted in **Figure 6a**, **6b**, and **6c**, with similar magnitudes. Notably, the GP for peaky speech and the ANM regressor for both stimulus conditions produced responses with much narrower and larger amplitude peaks (Wave V), suggesting superior temporal alignment with the subcortical potential’s fine structure. A figure with high-passed filter at 150 Hz applied to the ABRs that better highlights the early waves is provided in supplemental materials (**Figure S2**).

**Figure 6.**
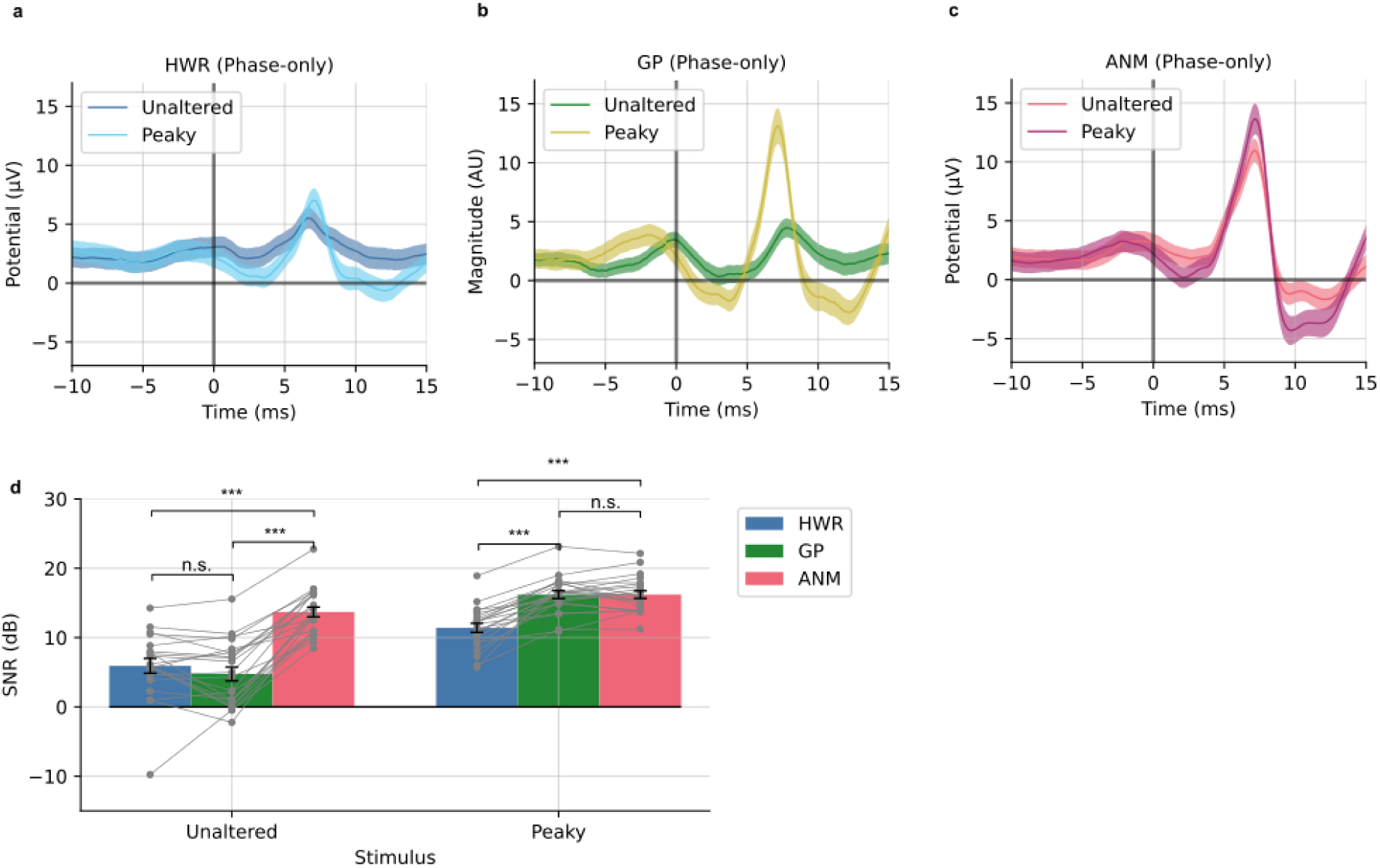
The ABR derived from the phase-only version of the three regressors: HWR **(a),** GP **(b),** ANM **(c)**. **(d)** The SNR analysis for the phase-only regressor-derived ABRs.

Finally, we performed an SNR analysis analogous to that performed with the original regressors. Again, we found the regressor effect on SNR outcomes in both unaltered and peaky speech conditions (p<0.001; repeated measure ANOVA; **Figure 6d**). Specifically, in unaltered speech, the ANM regressor surpassed both HWR and GP (p<0.001; two-tailed paired t test, Holm-Bonferroni corrected). In peaky speech, the ANM and GP exhibited comparable SNRs (p=0.97) with both significantly outperforming HWR (p<0.001; two-tailed paired t test, Holm-Bonferroni corrected). Moreover, we found a significant effect of stimulus condition, where unaltered speech had generally lower SNR than the peaky speech (p<0.001 for stimulus condition variable), but this may have been partially driven by the inclusion of the GP responses to unaltered stimuli.

## Discussion

This study presents a comprehensive quantitative analysis and comparison of deconvolution paradigms designed to derive the human ABR from continuous naturalistic speech. This deconvolution method generally has the advantages that allow for investigating subcortical encoding of speech in a more natural and more engaging settings. We analyzed EEG recordings from subjects listening to both unaltered speech and modified peaky speech. We compared three regressors used in a deconvolution paradigm that were developed in recent studies: the HWR from Maddox and Lee (2018), the GP from Polonenko and Maddox (2021b), and the ANM from Shan et al. (2024). Several metrics were conducted to compare these regressors’ performance, including the derived ABR waveform SNR, the time required for subject to get robust ABR, and the prediction accuracies of the ABR kernel with broadband (Pearson’s correlation) and per-frequency (spectral coherence) approaches. The insights gained from these evaluations are intended to inform and guide future research in selecting the most appropriate regressors for ABR derivation from continuous, naturalistic speech.

### Quantitatively comparing deconvolution paradigms

To generalize our results: we found that the ANM regressor for both peaky and unaltered speech and the GP regressor for peaky speech provided the best performance. Some caveats and specific situations where one technique might be favored over another are discussed below. The HWR regressor provided relatively poor ABRs for both speech conditions. We also derived the response from natural speech using the GP regressor for completeness, but we do not recommend this combination for practical use. Even though it did yield an ABR, the quality was predictably bad, with responses showing small amplitude and broad Wave V. This combination is not discussed further.

The HWR regressor, which was the first of these techniques to be developed (Maddox & Lee, 2018) did not match the performance of other regressors in either speech condition. The HWR-derived ABR exhibited a relatively noisy waveform with a broad Wave V (**Figure 2a**), requiring more than 42 minutes to acquire robust ABRs from all subjects (SNR>=0 dB; **Figure 3b**). The HWR ABR kernel resulted in low prediction accuracy because the kernel lacked the temporal detail of subcortical responses and had lower SNR. However, when extended the kernel time window to incorporate the response with cortical responses, its performance was similar to the other two regressors (**Figure 4a**).

The GP regressor coupled with peaky speech provided ABRs that showed early waves (Wave I) in the raw responses. When high-passed at 150 Hz, both Wave I and Wave III could be seen (**Figure S1**), allowing for examining the early generators of the auditory evoked potential. These responses may hold potential for clinical use. The GP regressor was also more efficient than HWR, with all subjects reaching the 0 dB SNR criterion in only 12 minutes. This efficiency could be further enhanced, as the prior study has shown, with high-pass filtering at 150 Hz potentially reducing the time to around 5 minutes (Polonenko & Maddox, 2021b). GP-derived kernels also provided better prediction than HWR.

The ANM regressor demonstrated superior performance in unaltered speech and comparable performance as GP in peaky speech conditions. This regressor did not only derive the best SNR ABR, but like the GP’s ability in peaky speech, this regressor also has the benefit of showing early ABR components—Wave I and Wave III—for both speech conditions, even without the needs of further filtering (**Figure 2c**). The time required to get decent ABR in both conditions was substantially reduced compared to HWR, and it was even faster than GP for peaky speech (**Figure 3b**). The best prediction accuracy was achieved using the ANM-derived kernels in both unaltered and peaky speech in correlation and spectral coherence analysis. The ANM’s excellent performance stems from its biological fidelity, as it takes the auditory system’s peripheral nonlinearities into account in the linear deconvolution paradigm, with the adaptation in the auditory nerve being particularly important (Kulasingham et al., 2024; Shan et al., 2024).

Some of the metrics we tested, such as SNR and acquisition time (as well as general waveform morphology) are frequency dependent, and thus affected by the power spectrum of the regressors, which differed substantially (**Figure 5**). Because deconvolution can be computed through frequency domain division, using spectrally different regressors is equivalent to applying different filtering to the EEG data (and equivalently, to the deconvolved response). These differences mean that direct comparison of the responses with different filtering might not be fair, because one regressor may accentuate noisier frequency bands than others. We attempted address this issue and provide a fairer comparison by setting the magnitude of each regressor to one across frequencies, preserving only the phase information. The deconvolved responses maintained a typical ABR morphology, where the GP for peaky and the ANM for both conditions showed about the same magnitude of wave V, both surpassing that of HWR. In the SNR analysis of these phase-only regressor derived ABRs, ANM once again led in performance for unaltered speech, with both GP and ANM outperforming HWR in peaky speech conditions, with no notable difference between the former two. These findings suggest that the ANM provides superior performance with unaltered speech, but with peaky speech either the ANM or GP can be used, with the choice depending more on experimental factors, as discussed below. With the phase-only regressors, better synchronization should result in larger waves in the deconvolved responses, which is confirmed in **Figure 6** for the GP and ANM regressors. The phase-only regressor was designed as an approach to compensate for spectral differences when comparing regressors—whether it provides value as an additional method for deriving responses will require further consideration.

Finally, we found that, between the two stimulus types, peaky speech elicited subcortical EEG responses that could be predicted with higher accuracy than unaltered speech. When analyzing phase-only regressors, this trend holds true across all regressors, with peaky speech resulting in superior SNR regardless of the regressor employed. Even the HWR-derived ABR from peaky speech had better SNR than unaltered speech. Similar to the CHEECH (CHirp-spEECH) stimuli (Backer, Kessler, Lawyer, Corina, & Miller, 2019) that incorporated chirps into speech, peaky speech is designed to make the speech click-like, aligning neural responses across the tonotopic axis, thereby eliciting stronger auditory evoked potentials (Polonenko & Maddox, 2021b). Given that peaky speech hardly alters sound quality and does not impact intelligibilty, it stands out as the preferable stimulus for deriving speech-evoked ABR, when experimental conditions allow.

### Qualitatively comparing deconvolution paradigms

The natural question following a comparison of two stimulus types and three regressors is what to use in future experiments. Since the ANM regressor (for both stimulus types) and GP regressor (for peaky speech) provided very similar performance, the answer is nuanced and experiment-dependent (the HWR regressor was poorest by all metrics and is unlikely to be appropriate). Both of these regressors and both stimulus types have their strengths and weaknesses that will determine the best choice.

Where the GP regressor is discussed below, it is on the assumption that peaky speech is used as the stimulus. The overall shape of the response is not considered a differentiating factor between the GP and ANM regressors because they can be made to be very similar through spectral manipulation (i.e., filtering).

We will discuss stimulus type first. Peaky speech’s primary disadvantages are that it requires pre-processing and that it is not quite natural, although we consider the latter issue to be minor. It also cannot be broadly applied to arbitrary stimulus types, as it assumes a calculable fundamental frequency. Its advantages are that it can be used with either the GP or ANM regressor, affording greater flexibility for analysis, and provides slightly better responses than natural speech with both regressors. Natural speech, beyond the obvious benefit of its inherent ecological validity, has the advantage of needing no pre-processing, making it appropriate for real-time use where sound and EEG data are recorded at the same time. Natural speech cannot be used with the GP regressor, so requires that the ANM be used for analysis (or similar methods, as described in Kulasingham et al. (2024)).

A unique benefit of the GP regressor is that the impulses that make up the pulse train regressor are of unit magnitude, meaning the deconvolved ABR can be expressed in meaningful units of electrical potential. While not explored in this study, Polonenko and Maddox (2021b) highlighted another benefit of using the GP regressor with multiband peaky speech, where the GP regressor can be extended to simultaneously investigate ABRs across different frequency regions, working on a similar principle to the parallel ABR (Polonenko & Maddox, 2019), offering a broader clinical application scope.

The ANM does not require pre-processed stimuli and is useful for studying a wide range of spectro-temporally rich natural stimuli, including music (Shan et al., 2024), making it versatile for various research purposes. However, compared to the GP, it has the limitation that the derived ABR is not expressed in meaningful units. Computing the ANM regressor takes considerable computation time, although this can be mitigated by using similar regressors that still include adaptation (Kulasingham et al., 2024). Thus, while the GP requires significant stimulus pre-processing, use of the ANM regressor requires substantial processing at the analysis stage. In the majority of use cases, neither of these requirements pose a problem, as stimulus and regressor generation are both typically one-time offline procedures. A recent study by Kulasingham et al. (2024) compared the ANM with regressors generated by other simpler auditory periphery models. They found that when using a more computationally efficient regressor that still includes nonlinear effect of adaptation (Osses Vecchi & Kohlrausch, 2021), the SNR of the derived ABR is similar to that of the more complicated ANM, despite the derived ABR’s lack of early components (Kulasingham et al., 2024).

A limitation of our study was its exclusive focus on a single speech stream narrated by a male speaker. Previous studies indicate that speech from a female speaker, characterized by a higher pitch, tends to reduce the amplitude of wave V (Polonenko & Maddox, 2021b; Saiz-Alía & Reichenbach, 2020). This effect is particularly relevant for peaky speech, where a higher pitch correlates with a faster rate, leading to neuronal adaptation and refractoriness (R. Burkard & Hecox, 1983; R. Burkard, Shi, & Hecox, 1990). Another potential issue in need of further study is how well the ANM works when separately deconvolving speech streams that were presented simultaneously. Primary indications from data recorded by our lab but not included here is that the ANM struggles in this setting where the GP does not. This concern is noted here because we consider it practically important, but it will need further exploration.

Finally, it is important to consider what the deconvolved response really represents. Calling it a response is a bit of a misnomer—it is a temporal kernel that relates a regressor to an EEG recording through convolution. This distinction is not pedantic. Consider an example experiment in which the same peaky speech stream is presented at a high and low level 20 dB apart. The subcortical response to the lower level stimulus will be smaller and later. The GP regressor is the same for both stimulus levels, and the deconvolved ABR should be smaller and later, as expected. The ANM regressor, however, changes based on stimulus level. The regressor itself will be smaller and later at the lower level. If we assume it perfectly estimates the change, then the deconvolved response will be the same for both stimulus levels. If the ANM overestimates the amplitude reduction and delay, then the deconvolved ABR could even be larger and earlier for the lower stimulus level, which would be a very strange result on its face.

It could still be that the ANM is the best regressor for that experiment, but here and generally, careful consideration must be given to the design, analysis, and interpretation of deconvolution studies.

## Materials and Methods

### EEG Dataset

The data analyzed in this study were obtained from a broadband peaky speech experiment previously conducted by Polonenko and Maddox (2021a). In that experiment, EEG was recorded from 22 normal hearing subjects (aged 18–32 years, mean ± SD of 23.0 ± 3.6 years) while they listened to the audiobook *The Alchemyst* (Scott, 2008), which was narrated by a male voice, as detailed in Polonenko and Maddox (2021b). The silent pauses exceeding 0.5 s in the audiobook had been truncated, and the audiobook was segmented to forty excerpts, each lasting 64 seconds. The recording time was 42 minutes and 40 seconds for each stimulus condition. During the experiment, subjects passively listened to the speech stimuli over ER-2 insert earphones at an average sound pressure level of 65 dB.

The EEG signal capturing subcortical activity (used to compute the ABR) was recorded using BrainVision’s passive Ag/AgCl electrodes. These electrodes were placed at the frontocentral position (FCz in the 10-20 system, active non-inverting), on the left and right earlobes (inverting references), and at the frontal pole (Fpz, ground). The electrodes were connected to an ActiCHamp system with the signal sampled at 10 kHz and high-pass filtered at 0.1 Hz. The recording process also applied a causal, fourth-order lowpass filter at 1/3 Nyquist (1667 Hz). Subsequent offline preprocessing included applying a high-pass filter at 1 Hz to remove any slow drifts, and a notch filter at 60 Hz along with its first three odd harmonics to reduce power line noise.

### Stimuli

In the original dataset, subjects listened to three stimulus conditions. However, for the purpose of this study, we focused on analyzing only two of those conditions: 1) unaltered speech, 2) re-synthesized broadband peaky speech. The re-synthesized peaky speech was designed to make the speech audio impulse-like by aligning the phase of the harmonics at the time of glottal pulses. This design aimed to elicit brainstem responses similar to those elicited by clicks, thereby evoking canonical ABRs while still preserving the intelligibility of the speech with minimal perceptible differences from the unaltered version. For a detailed explanation of the peaky speech synthesis process and audio examples, see Polonenko and Maddox (2021b).

### ABR Derivation

#### Deconvolution model for ABR

As described in Maddox and Lee (2018), Polonenko and Maddox (2021b), and Shan et al. (2024), an encoding model of the ABR was defined as shown in **Figure 1a**. The speech stimuli were processed differently to isolate a given stimulus feature (i.e., regressor) to be used as the input *x*, while the EEG signal was the output *y*, and the ABR was the impulse response of a linear system and determined through deconvolution. The computation was performed in the frequency domain for efficiency:

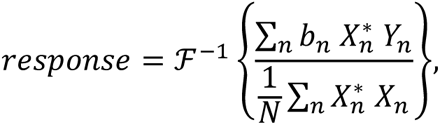

where *response* denotes the derived impulse response (i.e., the ABR), *X*_*n*_ the Fast Fourier transform (FFT) of the regressor for trial *n*, *Y*_*n*_ the FFT of EEG signal for trial *n*, * the complex conjugate, ℱ^−1^the inverse FFT, *b*_*n*_ the weight for trial *n* (see below), *N* the total number of trials, and *n* the trial index.

When computing the average response, a Bayesian-like process (Elberling & Wahlgreen, 1985) was used to account for variations in noise level, so that noisier trials were weighted less. The EEG recording from each trial was weighted by the inverse variance, 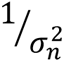, relative to the sum of the inverse variances of all

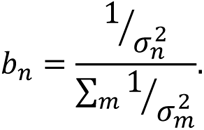

### Three Regressors

We compared the three regressors from previous three studies:

1) Half-wave rectified stimulus (HWR; **Figure 1b**) The half-wave rectified stimulus regressor was generated by first taking the positive values of the stimulus waveform and downsampling it to 10 kHz. This positive component of the stimulus was then used as the input to the encoding model (i.e., *x*), denoted as HWR. Then, the same process was applied, but with the original stimulus inverted so that the negative values (now positive) were used, and downsampled as before. Deconvolution was performed independently using both the positive and negative components as inputs. The final ABR response for each epoch and each subject was computed by averaging the responses to the positive and negative components.
2) Glottal Pulse (GP; **Figure 1c**) The glottal pulse times were initially extracted from the speech stimuli using speech processing software, *PRAAT* (Boersma, 2011) when the peaky speech stimuli were constructed. The sequence of impulses that occurred at the glottal pulse times in the peaky speech stimuli was then used as the input to the encoding model, denoted as GP.
3) Auditory Nerve Model firing rate (ANM; **Figure 1d**) A computational auditory periphery model created by Zilany, Bruce, Nelson, and Carney (2009), updated in Zilany, Bruce, and Carney (2014), and adapted for Python (Rudnicki, Schoppe, Isik, Völk, & Hemmert, 2015) was utilized to generate simulated auditory neural responses. It was previously shown to be able to account for the peripheral nonlinearity effects (Kulasingham et al., 2024; Shan et al., 2024). The speech stimuli were upsampled to 100 kHz according to the model’s requirement and converted to a pressure waveform (measured in pascals) at 65 dB SPL. We set the characteristic frequency (CF) ranging from 125 Hz to 16 kHz spaced at 1/6 octave intervals. The auditory nerve firing rate was then summed across all CFs of high spontaneous rate fibers and downsampled to match the EEG sampling rate of 10 kHz so it could be utilized as the regressor, denoted as ANM.

The three regressors were used for both the unaltered and the peaky speech conditions. Although the GP regressors were intended for use with peaky speech and have limited effectiveness as representative features for unaltered speech, they still capture some acoustic information on the timing of glottal pulses in unaltered speech.

### Phase-only Regressors

Since the three different regressors have different spectra as shown in **Figure 5**. Direct quantitative comparison on the time-domain response waveforms derived from these regressors (e.g., wave V amplitude) is not applicable, since they all have different spectra and overall magnitudes. Thus, we modified our analysis so that the derived response does not depend on the regressor magnitude spectrum by using phase-only regressors, generated as follows:

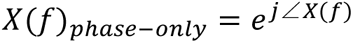

where *X*(*f*) is the FFT of the regressor and ∠ *X*(*f*) is the angle of the regressor in frequency domain.

The phase-only version of the three regressors were then used to derive ABRs using the deconvolution encoding model.

### Performance Metrics and Statistical Analysis

#### Response Signal-to-Noise Ratio (SNR)

To evaluate the quality of the derived ABRs, we estimated the SNRs of each waveform as described in previous studies (Maddox & Lee, 2018; Polonenko & Maddox, 2021b; Shan et al., 2024) using the following equation

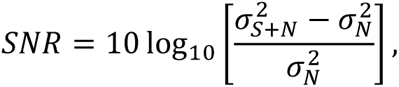

where 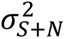 is the variance of the ABR waveform measured within the time interval of 0 to 15 ms, and 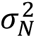 is the noise variance computed by averaging the variance across each non-overlapping 15 ms segment within the pre-stimulus baseline period, spanning from −200 to −20 ms. Therefore, subtracting 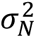 from 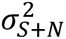 in the numerator offers an estimate of the signal variance 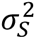, which is then divided by the noise variance and log transformed and scaled to estimate SNR in decibels.

SNR was analyzed through a repeated measures ANOVA followed by a post-hoc pairwise t-test to compare the three regressors. This comparison was also applied separately to the three phase-only regressors.

### Time Required to Obtain Robust Responses

We were interested in how long it took to record data in order to get a robust ABR using each of the three regressors. For each subject, we calculated the SNRs of ABRs, using the previously mentioned formula, across a recording duration ranging from 1 to 42 minutes. We then reported the cumulative proportion of subjects who achieved an ABR with an SNR of at least 0 dB throughout the recording process, as in the original peaky speech study (Polonenko & Maddox, 2021b).

### Correlation between the predicted and the real EEG

To compare the power of the regressors to predict EEG, we used the responses to predict the EEG and calculated the correlation coefficient between the predicted EEG and the real EEG data, as in our previous study (Shan et al., 2024). The predicted EEG were generated by utilizing the ABRs from each regressor as kernels (full kernel: [0, 200] ms time range; subcortical kernel: [0,15] ms time range), which were then convolved with the corresponding stimulus’s regressors. We then calculated the Pearson correlation coefficient between the predicted and real EEG data as a performance metric for each regressor.

### Spectral Coherence

The ability of the regressors to predict EEG across different frequencies was evaluated using spectral coherence analysis, as outlined in Shan et al. (2024). This approach served as a normalized correlation between the predicted EEG and the real EEG data but is split across various frequency bins. To determine spectral coherence, the predicted EEG and the real EEG data were sliced into segments of specific window sizes (0.2 s in this study), which then determined the frequency bins. The coherence of each of these frequency bins was computed as the following equation

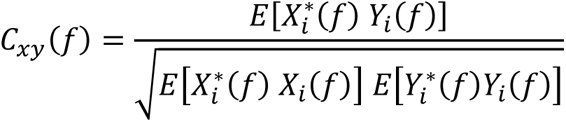

where *C*_*xy*_(*f*) denotes the coherence between signal *x* and *y* at frequency bin *f*, *E*[] is the expected value across slices, * the complex conjugate, *X*_*i*_(*f*) the FFT for predicted EEG slice *i* in frequency bin *f*, and *Y*_*i*_(*f*) the FFT for real EEG data slice *i* in frequency bin *f*.

To estimate the noise floor of the spectral coherence, we shuffled the order of the predicted EEG and real EEG data and calculated the spectral coherence for these mismatched trials. The median coherence value from these mismatched trials served as the noise floor shown in **Figure 4c** and **4d**.

To compare the performance of the three regressors in spectral coherence analysis, we computed the mean of the absolute value of the spectral coherence across three specific frequency bands for each regressor. These three frequency bands — [0, 25] Hz, [25, 85] Hz and [85, 135] Hz were selected based on findings from a previous study indicating superior performance of ANM in these ranges compared to HWR.

### Statistical Test

Data were checked and confirmed for normality using the Shapiro–Wilk test in parametric test. To compare the performance metrics of the three regressors, mixed effects linear regression models were constructed (formula below) in python, using stimulus condition, regressor and their interaction as the fixed effects and subject as random effect.

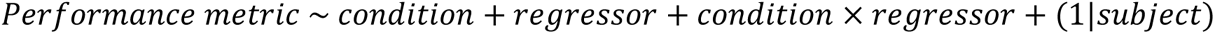

To compare the metrics within each stimulus condition, repeated measure ANOVAs followed by a pairwise post-hoc paired t-test with Holm-Bonferroni correction were used. The performance metrics used in these statistical tests were SNR analysis, broadband prediction accuracy (Pearson correlation), and the mean absolute value of the spectral coherence from the three frequency bands, as described above.

## Notes

### Competing Interest Statement

The authors have declared no competing interest.

