## supplemental materials for "Comparing Methods for Deriving the Auditory Brainstem Response to Continuous Speech in Human Listeners"

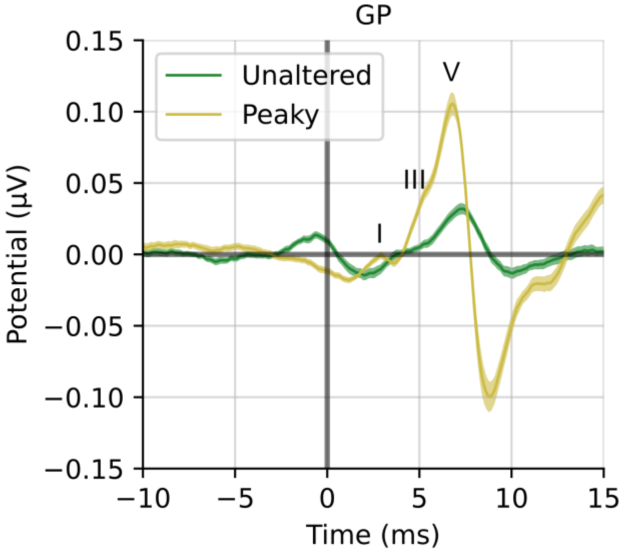


**Figure S1**. GP-derived ABR with high-pass filtered at 150 Hz to highlight the early waves. Wave I, III and V are annotated.

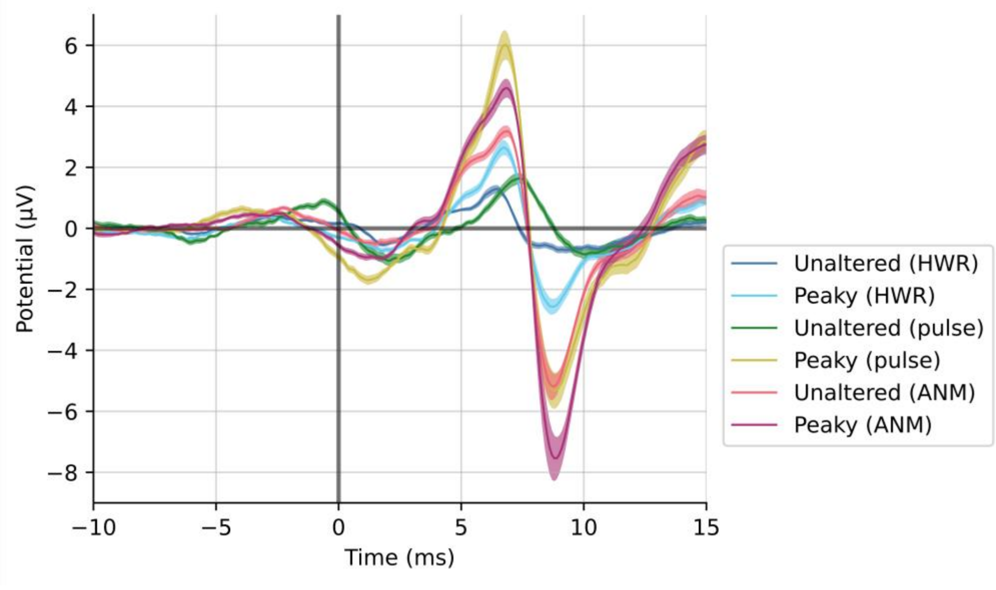


**Figure S2.** The ABRs derived from the three phase-only regressors with high-pass filtered at 150 Hz.
